# Artery-Like Smooth Muscle Drives Contractile Function in Dural Venous Sinuses

**DOI:** 10.64898/2026.06.01.729135

**Authors:** Madison E. Lemire, Hannah C. Ryan, Lillian S. Hand, John C. DeWitt, Liqun He, Christer Betsholtz, Adam S. Sprouse Blum, Nicholas R. Klug

## Abstract

The dural venous sinuses are the main conduit for cerebral blood flow to drain from the head back to the heart. Pathology of these sinuses has significant impacts on cerebral blood volume and intracranial pressure, indicating that these veins have an important role in cerebral hemodynamics. Active and modifiable mechanisms of sinus diameter control may present an opportunity for physiological or pathological regulation of venous blood flow. Here, we characterized the molecular and anatomical properties of dural vascular smooth muscle cells (SMCs) using RNA-sequencing and immunohistochemistry, demonstrating that artery-like SMCs are exclusive to sinus vessels. An ex vivo pressurized sinus preparation from mice was used to functionally characterize the vasodynamics of the sinus and its corresponding bridging veins. We found that the sinus contains dynamically contractile SMCs and constricts to both pressure and contractile agonists, while the bridging veins do not. We further demonstrated that these sinus SMCs exhibit dynamic calcium signaling that is responsive to contractile stimuli. These results reveal a contractile SMC phenotype localized to dural venous sinuses and demonstrate that sinus vessels possess active, artery-like mechanisms for regulating cerebral venous outflow.

## INTRODUCTION

The brain’s considerable metabolic demands are supported by a vast, highly organized vascular network responsible for delivering and draining blood. Sustained cerebral blood flow requires substantial pressure gradients that drive blood from the arterial system, through capillary beds, and ultimately into the venous circulation. The anatomical organization and flow capacity of these valveless veins vary widely, from small parenchymal vessels to large collecting venules, to dural venous sinuses, embedded within the outermost meningeal layer, the dura mater [1–4].

Aptly translated as “tough mother”, the dura mater is a collagen-rich, highly innervated fibrous tissue. Narrowing of the dural venous network is generally presumed to result from extrinsic factors, including shifting intracranial pressure, swelling of perisinus structures such as arachnoid granulations, or chronic intrinsic factors including pathological vascular remodeling that reduces luminal diameter [5, 6]. In contrast, active regulation of vascular tone is typically attributed to inlet vessels such as arteries and arterioles.

In both humans and rodents, dural venous sinuses experience substantial and dynamic intraluminal pressures around 3-10 mmHg in healthy conditions and rising above 20 mmHg in disease [7–11]. These pressures highlight their potential to influence cerebral venous outflow resistance and cerebrospinal fluid (CSF) drainage. Indeed, transient or chronic narrowing (stenosis) of venous sinuses produces robust alterations in blood and CSF drainage [12, 13]. Moreover, rapid re-stenosis frequently develops adjacent to stented sinus segments [14], suggesting that sinus caliber may be dynamically regulated rather than determined solely by fixed structural narrowing. Consistent with this possibility, recent in vivo observations demonstrate that the superior sagittal sinus undergoes rapid diameter changes during behavior-linked fluctuations in intracranial pressure [15], and that calcitonin-gene related peptide (CGRP) signaling can modulate sinus diameter over slower timescales [16]. However, the smooth muscle cell (SMC)-specific mechanisms that regulate sinus diameter under pathological and physiological conditions remain obscure. Other organ systems offer examples of dynamic venous contractility including those of the mesentery, lung, and liver [17–20]. Whether similar SMC-driven contractile features exist within the dura mater, and whether they influence cerebral venous drainage, remains unresolved.

Here, we directly assessed the molecular, anatomical, and functional contractile features of the cerebral venous network, with specific focus on dural bridging veins and venous sinuses. Using single-cell RNA sequencing, we characterized mural cell transcriptomic identities within the dura mater and highlight a unique mural cell population expressing markers of both venous and arterial identity. Whole-mount immunohistochemistry in mouse dura revealed that this venous–arterial hybrid mural cell population corresponds to SMCs confined specifically to large sinus vessels, absent from bridging veins and from the wider brain and dura vasculature. This anatomical pattern was also confirmed in human dural tissue. We then assessed the functional capacity of these venous SMCs in live, isolated, intact sinuses. Traditional contractile stimuli, including increased transmural pressure, membrane depolarization, and Gq-coupled receptor activation, evoked robust Ca^2+^ elevations and sinus constriction. Conversely, activation of K_ATP_ channels and nitric oxide signaling produced strong venodilation.

Together, these data demonstrate that contractile, artery-like properties within the cerebral venous system are uniquely concentrated in dural venous sinuses. These findings highlight the dynamic and contractile nature of sinus SMCs and expand our understanding of venous contributions to cerebral vascular dynamics, with potential implications for regulation of cerebral blood flow, CSF clearance, and intracranial pressure.

## RESULTS

### Single-cell RNA sequencing reveals distinct vascular smooth muscle cell populations including a population with a combined venous and contractile molecular identity

To define the transcriptomic identity of dural mural cells, we reanalyzed the mural cell population from a previously published single-cell RNA-sequencing dataset containing all isolated and sequenced cells from mouse dura mater [21] (Fig. 1A). Subclustering of these mural cells revealed three distinct smooth muscle cell populations and a population of pericytes (Fig. 1B). There was strong expression of canonical arterial SMC markers in the SMC1 population, including Pln, Acta2, and Sorbs2 [22] (Fig. 1B). The SMC2 population was likely venous SMCs, given the high expression of canonical venous SMC markers such as Ccl19 and Vtn and low expression of typical contractile markers such as Acta2. The SMC3 population exhibited overlap with markers from both SMC1 and SMC2, including high expression of both Ccl19 and Acta2, highlighting a shared venous and contractile molecular signature (Fig. 1B). These data identify a dural mural cell population with combined venous and contractile SMC features.

**Figure 1.**
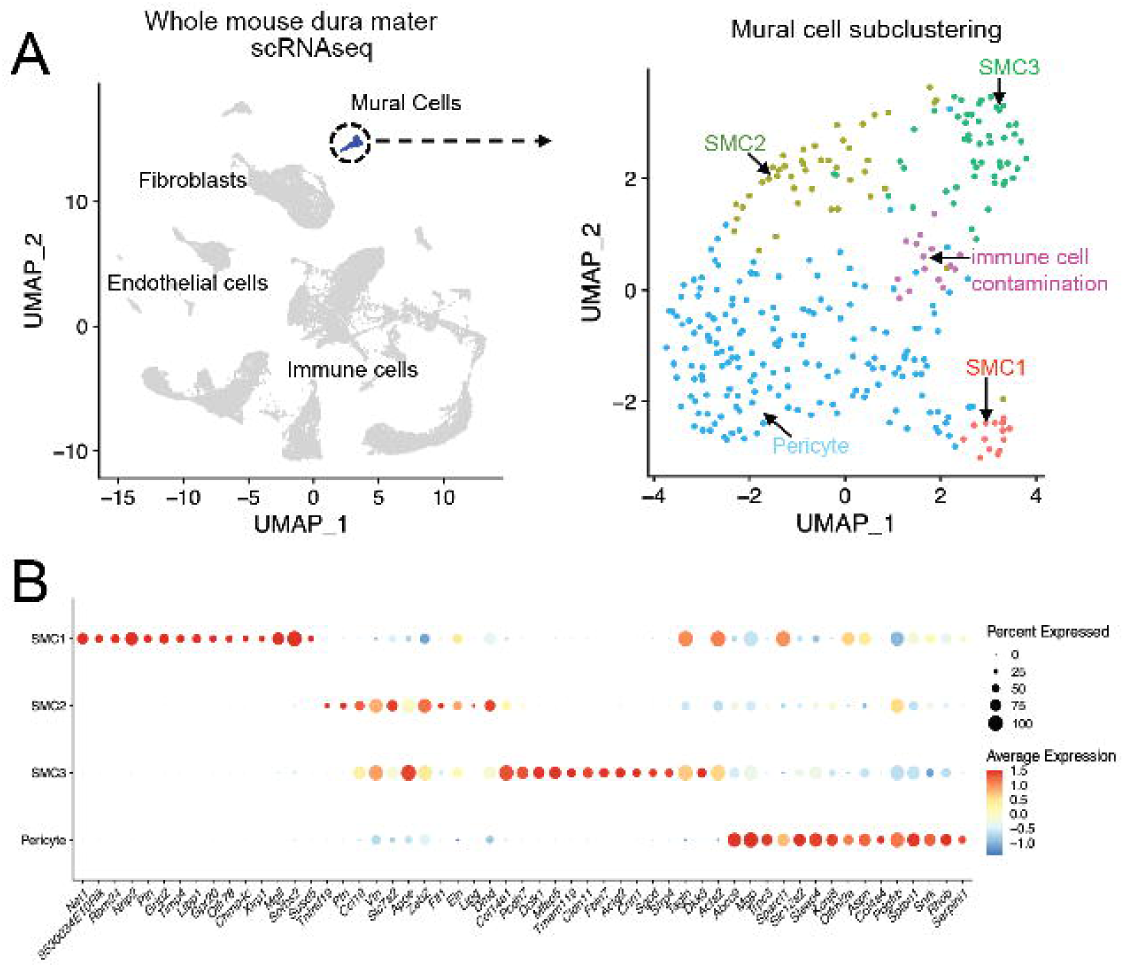
Mouse dura mater single-cell RNA-sequencing data identifies molecularly distinct mural cell populations. (A) Mural cells were extracted from a previously published single-cell RNA-sequencing dataset of adult mouse dura mater (Pietilä et al., 2023) and reanalyzed as a separate population. The original dataset is shown on the left, with the mural cell compartment highlighted for downstream analysis. Subclustering of extracted mural cells is shown on the right and resolved three smooth muscle cell (SMC) populations, designated SMC1, SMC2, and SMC3, along with a pericyte population and a small immune contaminant cluster. (B) Dot plot showing marker genes enriched across the mural cell subclusters. SMC1 is enriched for canonical contractile and arterial SMC-associated genes, including Pln, Acta2, and Sorbs2. SMC2 is enriched for venous-associated genes, including Ccl19 and Vtn, with relatively low expression of contractile SMC markers. SMC3 exhibits a combined venous and contractile molecular profile, including expression of both Ccl19 and Acta2. The pericyte cluster is distinguished by canonical pericyte-associated markers. Dot size represents the proportion of cells within each cluster expressing a given gene, whereas dot color denotes scaled average expression level.

### Dural venous sinus vessels contain smooth muscle cells with arterial contractile proteins that are absent in bridging and cerebral vein smooth muscle cells

To visualize protein expression and determine anatomical location of SMCs with shared venous and arterial molecular identity, we utilized immunohistochemistry on whole mount fixed dura mater tissue from mice. By staining with endothelial marker CD31 (PECAM1), a robust vascular network across the entire dissected dura mater tissue is visualized (Fig 2A). Apparent in this network are large bridging veins, which are severed (due to brain removal) at arachnoid bridging segments [23] and connect to transverse and superior sagittal sinuses (Fig. 2B).

**Figure 2.**
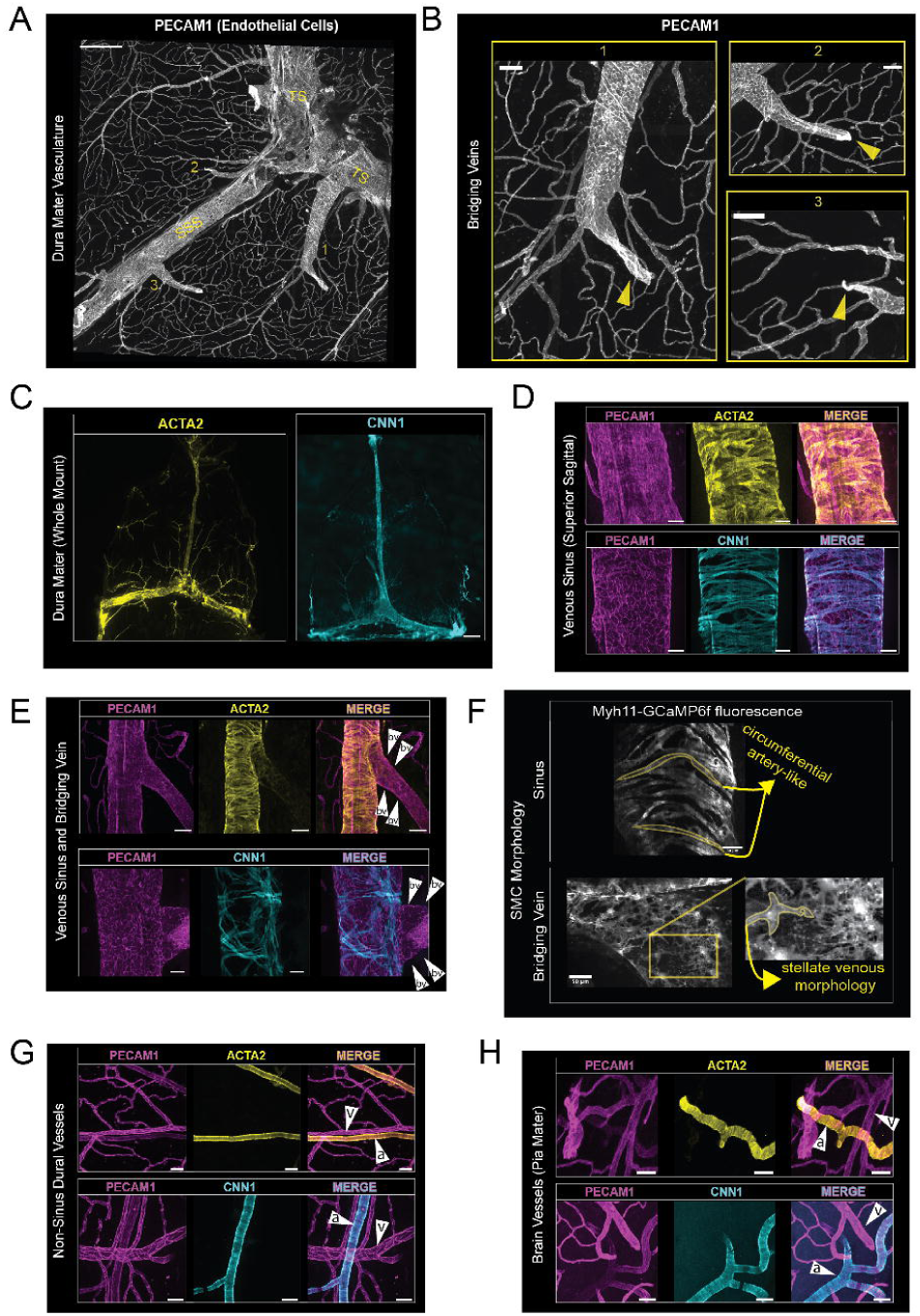
Contractile smooth muscle cell markers and artery-like morphology are restricted to dural venous sinuses. (A) Whole-mount mouse dura mater immunostained for the endothelial marker PECAM1/CD31 to visualize the dural vascular network. Stitched serial confocal z-stack imaging shows the superior sagittal sinus (SSS), transverse sinuses (TS), and associated bridging veins. Numbered regions indicate three bridging veins shown at higher magnification in panel B. (B) Higher-magnification images of the numbered bridging vein regions from panel A. Yellow arrows indicate cut bridging vein segments generated during tissue isolation, where bridging veins were separated from their arachnoid/pial vascular connections. (C) Whole-mount mouse dura mater immunostained for the contractile smooth muscle cell markers ACTA2/smooth muscle α-actin and CNN1/calponin-1. Widefield fluorescence imaging shows strong ACTA2 and CNN1 labeling along the dural venous sinuses. (D) Confocal z-projection of the superior sagittal sinus immunostained for PECAM1, ACTA2, and CNN1. ACTA2- and CNN1-positive smooth muscle cells are arranged along the sinus wall. (E) Confocal z-projection of a sinus–bridging vein junction immunostained for PECAM1, ACTA2, and CNN1. Contractile smooth muscle marker expression is prominent along the sinus and is sharply reduced or absent in the attached bridging vein. (F) Confocal z-projection of sinus and bridging vein mural cells from Myh11-GCaMP6f mice. Annotated cell borders highlight the elongated, circumferential morphology of sinus smooth muscle cells compared with the distinct stellate morphology of mural cells on the neighboring bridging vein. (G) Confocal z-projection of non-sinus dura mater vasculature immunostained for PECAM1, ACTA2, and CNN1. Contractile smooth muscle marker expression is observed on dural arteries/arterioles but is absent from nearby non-sinus dural veins. (H) Confocal z-projection of pial brain vasculature immunostained for PECAM1, ACTA2, and CNN1. Contractile smooth muscle marker expression is present on pial arteries/arterioles and absent from pial veins.

When whole-mount dura mater was stained for canonical arterial SMC markers, smooth muscle α-actin (ACTA2) and calponin-1 (CNN1), strong SMC labeling was observed specifically along the superior sagittal sinus and transverse sinuses, in addition to broad arterial labeling with ACTA2 and sparser labeling of arterioles with CNN1 (Fig. 2C). The sparse labeling of arterioles with CCN1 may explain the low population of *Cnn1* expressing SMCs seen in the arteriole SMC1 population (Fig. 1B). Confocal microscopy reveals long spindle shaped SMCs that express ACTA2 and CNN1 abundantly adorn sinus vessels and expression of these contractile proteins is rapidly lost at the sinus-bridging vein transition (Fig. 2 D, E). The artery-like SMC morphology found on sinus vessels also rapidly transitions to a venous-like stellate morphology [24, 25] at the bridging vein (Fig. 2F).

Outside of sinus regions, arteriole SMCs display prominent ACTA2 staining (in all arterioles) and CNN1 staining (in the sparser population of arterioles with CNN1 staining), whereas neighboring non-sinus veins lack smooth muscle cells expressing either marker (Fig. 2G). The artery SMC specific expression of ACTA2 and CNN1 was also observed in pial brain vasculature (Fig. 2 H). Thus, artery-like contractile protein expression is selectively enriched in dural venous sinus vessels and is absent from neighboring bridging and cerebral veins. We next asked whether this sinus-restricted SMC phenotype is conserved in human dura mater.

### Contractile phenotype of venous SMCs is confined to the sinus in human dura mater

Using postmortem formalin-fixed human dura mater samples we evaluated the sinus specific expression of contractile SMCs with ACTA2 staining. Coronal sectioning of human dura mater tissue allowed for clear identification of superior sagittal sinus vessel (Fig. 3A). Bridging veins were identified in serial slices with confirmation of merging lumen structures with superior sagittal sinus. Smooth muscle cells surrounding the superior sagittal sinus had prominent ACTA2 staining (Fig. 3B), a feature shared with dura mater arterioles (Fig. 3C). Bridging vein SMCs had absent ACTA2 staining (Fig. 3C) consistent with the loss of ACTA2 and other contractile marker staining in the bridging veins observed in mouse tissue (Fig. 2E). These human tissue data support the conclusion that ACTA2-positive SMCs are enriched along dural venous sinuses and reduced or absent in associated bridging veins.

**Figure 3.**
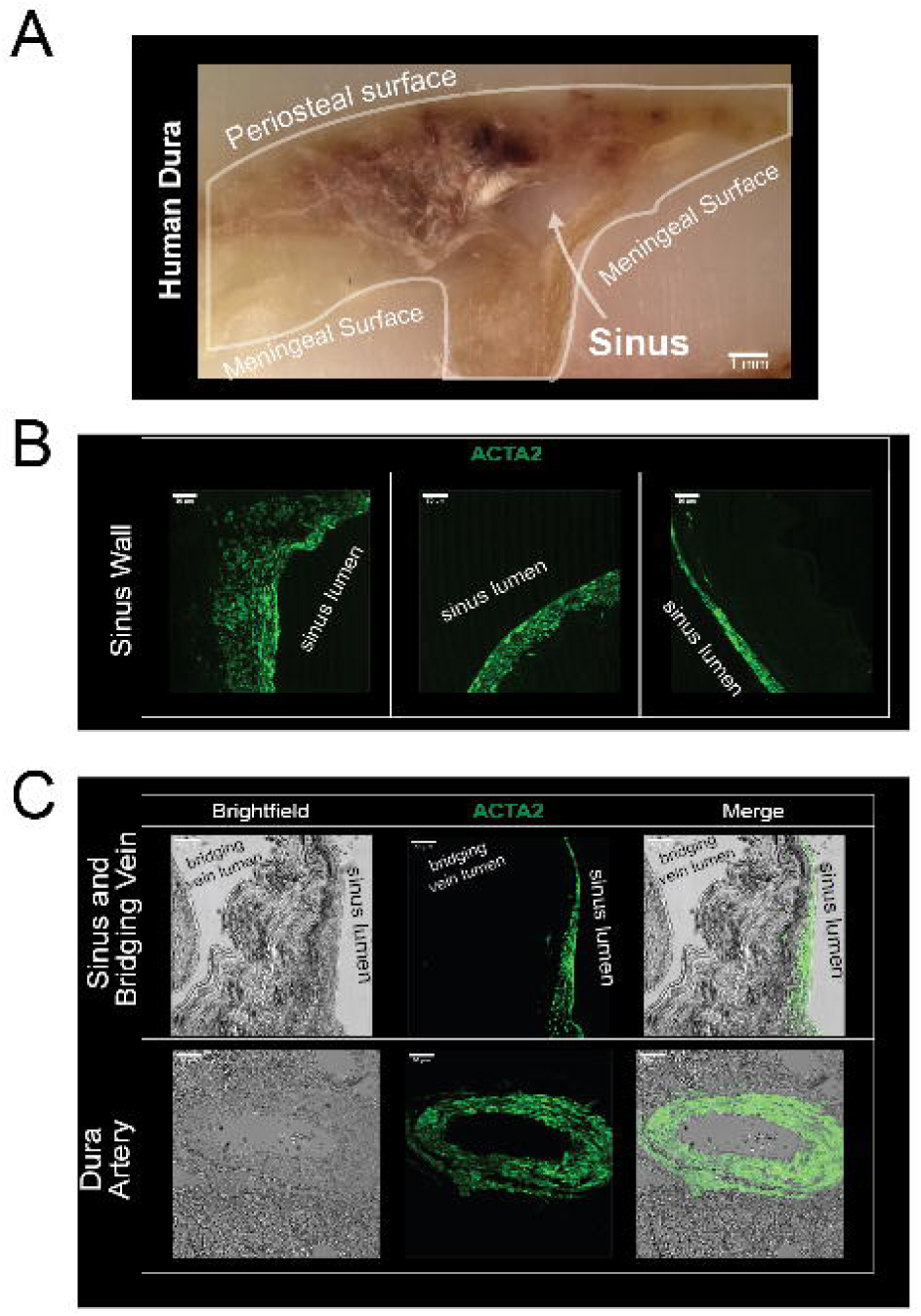
ACTA2 expression in human dural venous sinus vessels. (A) Coronal section of paraffin-embedded, formalin-fixed human dura mater containing the superior sagittal sinus. The periosteal surface, meningeal surface, and sinus lumen are annotated to show tissue orientation. (B) Higher-magnification confocal z-projections from multiple regions of the human sinus wall immunostained for smooth muscle α-actin (ACTA2), showing ACTA2-positive smooth muscle cells along the sinus wall. (C) Brightfield and confocal z-projection images of human dura mater regions containing the dural venous sinus, an associated bridging vein, and a dural artery. ACTA2 immunostaining highlights smooth muscle cells associated with the sinus wall and dural artery, with comparatively little ACTA2 labeling in the associated bridging vein. Scale bars: 1 mm in A; 50 µm in B and C.

### Sinus smooth muscle cells exhibit distinct Ca^2+^ events that are modulated by increased intraluminal pressure

Due to the anatomical and contractile protein similarities of sinus SMCs and artery SMCs we investigated whether live sinus SMCs exhibit Ca^2+^signaling responses typically seen in arterial SMCs. Specifically, the frequency of localized Ca^2+^ events in artery SMCs is coupled to intraluminal pressure, exerting a rapid intrinsic signaling feedback to regulate artery diameter [26, 27]. To determine whether the contractile SMCs surrounding dural venous sinuses were sensitive to changes in transmural pressure we isolated live whole mount dura mater preparations with physiological bath conditions and cannulated the superior sagittal sinus with an attached gravity column to change intraluminal pressure to deliver quantified transmural pressures (Fig. 4A). Using spinning-disk confocal microscopy, live tissue from myh11-GCaMP6f mice had robust expression of GCaMP6f in dural SMCs (Fig 4A). At 0 mmHg sinus SMCs generated abundant subcellular Ca^2+^ events (Fig. 4B, Movie 1). When exposed to increased intraluminal pressure, 5 mmHg sustained for at least 10 minutes, sinus SMCs exhibited an increase in Ca^2+^ event frequency, amplitude, and duration (Fig. 4C). These results demonstrate that sinus SMCs are not only anatomically contractile in appearance but also exhibit pressure-sensitive Ca^2+^ signaling. We next asked whether this pressure-sensitive signaling is accompanied by active changes in sinus diameter.

**Figure 4.**
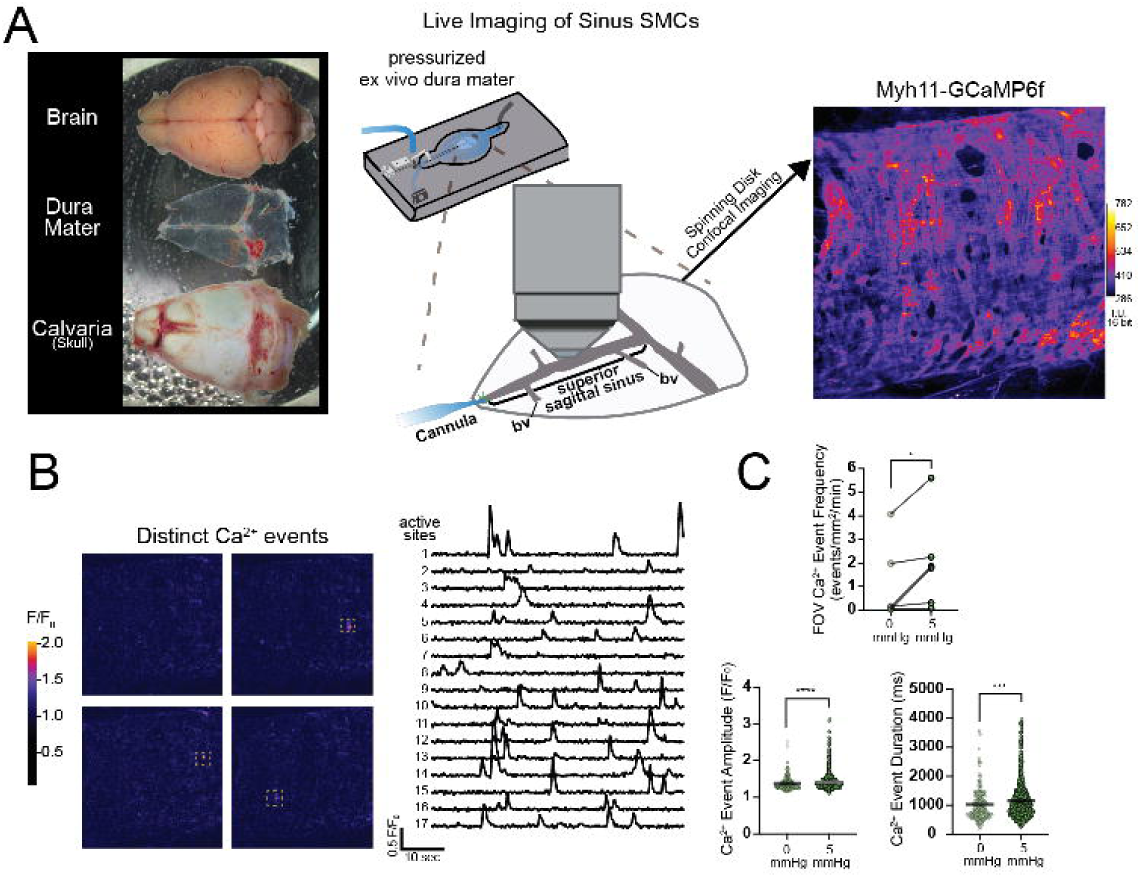
Intraluminal pressure increases localized Ca^2+^ event activity in dural venous sinus smooth muscle cells. (A) Live whole-mount dura mater preparation used to image Ca^2+^ activity in sinus smooth muscle cells. The dura was isolated from between the calvaria and brain, pinned in a bath-perfused recording chamber, and cannulated to allow controlled pressurization of the superior sagittal sinus. Schematic illustrates the ex vivo pressurized sinus preparation. Representative image shows raw pseudocolored GCaMP6f fluorescence in sinus smooth muscle cells from Myh11-GCaMP6f dural tissue. (B) Representative detection and analysis of localized Ca^2+^ events in sinus smooth muscle cells. Ca^2+^ events were extracted from raw GCaMP6f fluorescence using event detection criteria described in the Methods. Representative F/F_0_ traces show Ca^2+^ signals from 17 distinct subcellular sites within the field of view. (C) Quantification of localized Ca^2+^ event properties in sinus smooth muscle cells under unpressurized and pressurized conditions. Ca^2+^ event frequency, amplitude, and duration were measured across the field of view at 0 mmHg and after exposure of the same sinus smooth muscle cells to 5 mmHg intraluminal pressure. Individual values are shown with mean ± SEM; n = 6 preparations; n = 202 and 1441 individual events observed at 0 mmHg and 5 mmHg, respectively. *p < 0.05, **p < 0.01, ***p < 0.005, ****p < 0.001 by Mann–Whitney test.

### Dural venous sinus vessels constrict to intraluminal pressure

Blood and lymphatic vessels with contractile SMCs exhibit intrinsic contractile behavior in response to physiological levels of transmural pressure, where increased pressure leads to vessel narrowing to regulate perfusion [28, 29]. Using the ex vivo pressurized sagittal sinus preparation with near infrared microscopy we observed sharp contrast at the edges of superior sagittal sinus, allowing for real time diameter measurements of the pressurized sinus vessel (Fig. 5A). Immediately following pressurization (0 to 5 mmHg) the superior sagittal sinus exhibited overt constriction, with no observed change in bridging vein diameter (Fig. 5B, Movie 2). We further characterized the pressure-induced constriction response in the superior sagittal sinus and bridging vein vessels, by implementing a step-wise increases in intraluminal pressure from 1 to 20 mmHg. Superior sagittal sinus vessels demonstrated enhanced pressure-induced constriction compared to bridging veins at all pressure steps from 1 to 20 mmHg, and peak sinus constriction (39.3 ± 3.4%) was observed at 5 mmHg (Fig. 5C). Absolute diameter measurements of the superior sagittal sinus in 2 mM and 0 mM external Ca^2+^ highlight the robust Ca^2+^- sensitive constriction responses of the sinus SMCs (Fig. 5D). Together, these data demonstrate that dural venous sinuses, but not bridging veins, develop active Ca^2+^-sensitive constriction in response to increased intraluminal pressure. This pressure-dependent contractility suggested that sinus SMCs may share broader excitability features with arterial SMCs, including responsiveness to depolarization and receptor-mediated contractile stimuli.

**Figure 5.**
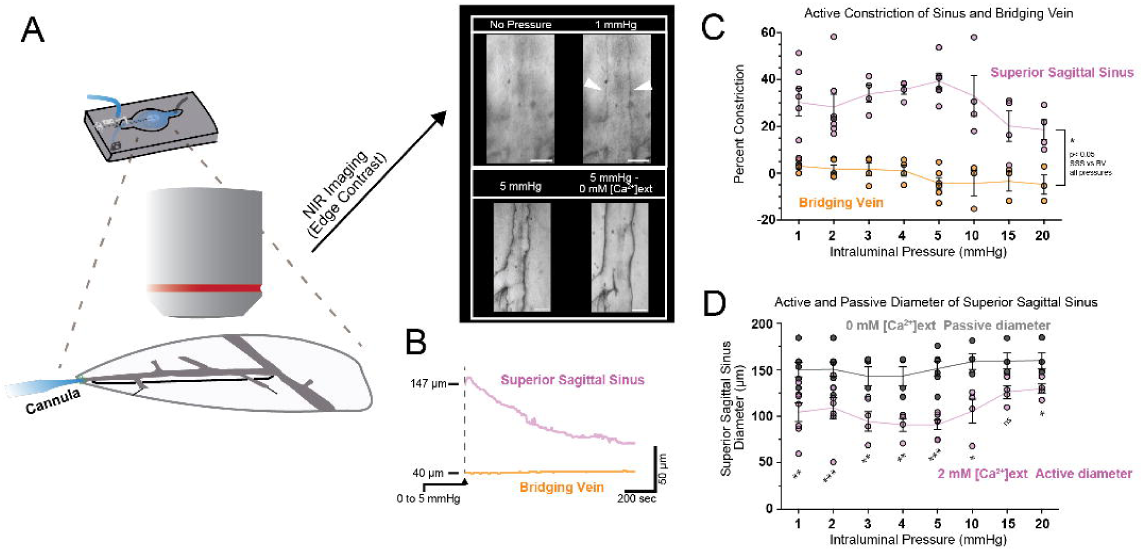
Dural venous sinuses constrict in response to increased intraluminal pressure. (A) Ex vivo pressurized sinus preparation used to assess pressure-induced changes in vessel diameter. Illustration shows the isolated, bath-perfused dura mater preparation with the superior sagittal sinus cannulated for controlled intraluminal pressurization. Representative infrared-differential interference contrast (IR-DIC) images show the superior sagittal sinus and attached bridging veins at 1 mmHg and 5 mmHg in normal physiological saline containing 2 mM extracellular Ca^2+^, and at 5 mmHg in Ca^2+^-free solution containing 0 mM Ca^2+^ and 5 mM EGTA. Vessel walls are visible by IR-DIC imaging, allowing edge-to-edge diameter measurements. (B) Representative diameter traces from the superior sagittal sinus and an attached bridging vein during a pressure step from 0 to 5 mmHg. The superior sagittal sinus constricts after pressure elevation, whereas the bridging vein shows little change in diameter. (C) Quantification of pressure-induced constriction in the superior sagittal sinus and bridging veins across intraluminal pressures from 1 to 20 mmHg. Percent constriction was calculated from active and passive diameters using the equation: ((passive diameter – active diameter) / passive diameter) × 100. (D) Absolute diameter measurements of the superior sagittal sinus in normal extracellular Ca^2+^ solution (2 mM Ca^2+^) and Ca^2+^-free solution (0 mM Ca^2+^ with 5 mM EGTA), demonstrating Ca^2+^-sensitive active tone in the pressurized sinus. Individual values are shown with mean ± SEM; n = 3–7 preparations. *p < 0.05, **p < 0.01, ***p < 0.001 by two-way ANOVA. Scale bar: 100 µm in A.

### Sinus smooth muscle cells exhibit global Ca^2+^ increases in response to mechanical, pharmacological, and sympathetic stimuli

Since intraluminal pressure modulated discrete Ca^2+^ events and caused robust constriction of the superior sagittal sinus, we utilized myh11-GCaMP6f tissue to assess whether pressure and other contractile stimuli typically attributed to arterial SMC excitability would elicit global increases in sinus SMC Ca^2+^ levels. Here, we examined the spatially averaged GCaMP6f fluorescence levels within all SMCs in the field of view (FOV) before and during each stimulus condition. In response to pressure (5 mmHg), depolarization (60 mM external K^+^), and thromboxane A2 receptor stimulation (1 uM U46619), sinus SMCs had robust and reliable increases in global Ca^2+^ levels (Fig. 6A,B). Movies 3, 4, and 5 demonstrate the sinus SMC Ca^2+^ response to pressure, depolarization, and thromboxane A2 receptor stimulation, respectively. Consistent with previous observations [30], the superior sagittal sinus is richly innervated with norepinephrine containing sympathetic nerve fibers (Fig. 6C), therefore we tested whether sinus SMCs respond to pharmacological α_1_ receptor stimulation with phenylephrine. Indeed, activation of α_1_ adrenergic receptors elicited a large global increase in SMC Ca^2+^ levels (Fig. 6D,E, Movie 6). Further, we also observed a robust increase in sinus SMC global Ca^2+^ in response to sympathetic nerve activation with stimulating electrodes (electric field stimulation, EFS) (Movie 7), a response that was strongly reduced by preincubation with the α_1_ adrenergic receptor antagonist, prazosin (Fig. 6F). Together, these data show that sinus SMCs respond to pressure, depolarization, Gq-coupled receptor activation, and sympathetic nerve stimulation with global Ca^2+^ elevations.

**Figure 6.**
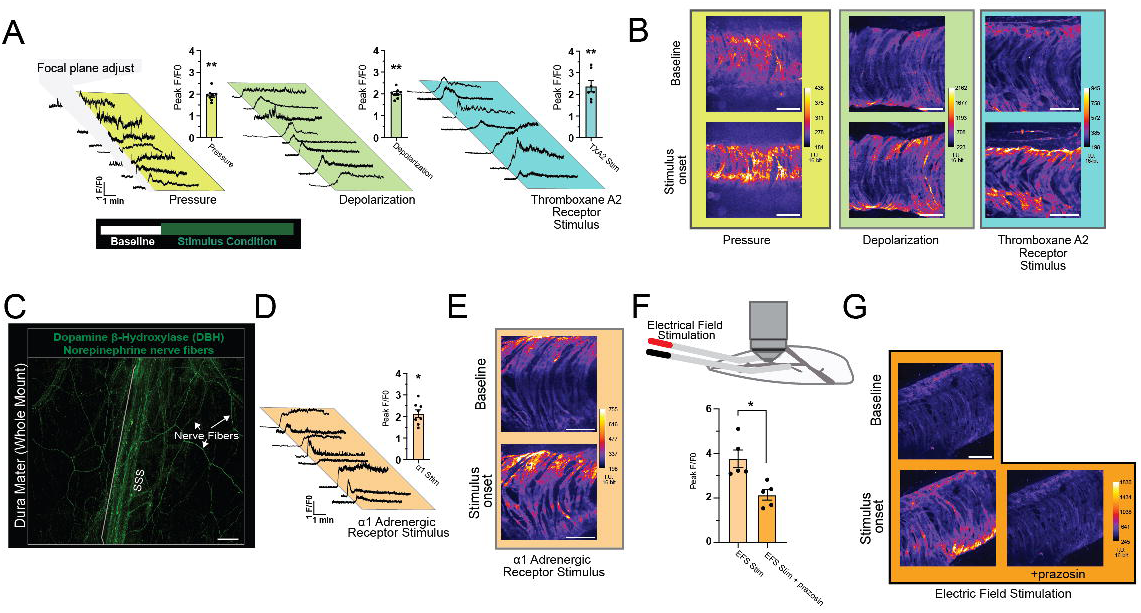
Contractile stimuli and sympathetic activation evoke global Ca^2+^ elevations in dural venous sinus smooth muscle cells. (A) Global cytosolic Ca^2+^ responses in sinus smooth muscle cells measured from Myh11-GCaMP6f dural tissue. Fluorescence changes were quantified as F/F_0_ across all in-focus sinus smooth muscle cells within the field of view, distinct from the localized subcellular Ca^2+^ event analysis shown in Figure 4. Representative F/F_0_ traces and corresponding peak F/F_0_ responses are shown before and after stimulation with increased intraluminal pressure from 0 to 5 mmHg, membrane depolarization with 60 mM extracellular K⁺, and thromboxane A2 receptor activation with 1 µM U46619. (B) Representative raw pseudocolored GCaMP6f fluorescence images from sinus smooth muscle cells before and after stimulation with pressure, 60 mM extracellular K⁺, and U46619. (C) Confocal image of the superior sagittal sinus and surrounding dural tissue immunostained for dopamine β-hydroxylase (DBH), showing dense DBH-positive sympathetic innervation along the superior sagittal sinus. (D) Representative F/F_0_ trace from sinus smooth muscle cells before and after α_1_-adrenergic receptor stimulation with 100 µM phenylephrine. (E) Representative raw pseudocolored GCaMP6f fluorescence images from sinus smooth muscle cells before and after α_1_-adrenergic receptor stimulation with 100 µM phenylephrine. (F) Peak global Ca^2+^ responses in sinus smooth muscle cells evoked by electrical field stimulation (EFS) of perivascular nerves. Platinum electrodes were positioned along the sinus border, and EFS was delivered at 10–90 V, 0.25 ms pulse duration, and 20 Hz. Responses are shown under control conditions and after bath application of the α_1_-adrenergic receptor antagonist prazosin (1 µM). (G) Representative raw pseudocolored GCaMP6f fluorescence images from sinus smooth muscle cells before and after EFS under control conditions and after EFS in the presence of prazosin. Individual values are shown with mean ± SEM; n = 4–8 preparations. For panels A and D, statistical significance was determined by Wilcoxon signed-rank test versus a hypothetical value of zero, representing no increase in Ca^2+^ signal. For panel F, statistical significance was determined by Mann–Whitney test. *p < 0.05, **p < 0.01, ***p < 0.005, ****p < 0.001.

### Smooth muscle cells regulate sinus diameter in response to contractile and relaxation stimuli

Lastly, we utilized the pressurized sinus preparation to assess the influence of sinus SMC excitability on vessel diameter. Application of isotonic 60 mM K^+^ bath solutions to depolarize SMCs generated a 63.4 ± 4.5% constriction in the sinus (Fig. 7A). Stimulation of SMC α_1_ adrenergic receptors with 100 uM phenylephrine, and thromboxane A2 receptors with 1 uM U46619 produced 41.8 ± 5.8% and 56.5 ± 3.2% constriction, respectively, in the sinus (Fig. 7 B,C). In contrast to sinus vessels, bridging veins constricted a modest, yet significant 3.6 ± 1.1% in 60 mM K^+^ conditions. Bridging vein constriction was undetected in phenylephrine and U46619 conditions (Fig. 7 B-C). Movies 8, 9, 10 visualize sinus contractile responses to 60 mM K^+^, phenylephrine, and U46619, respectively.

**Figure 7.**
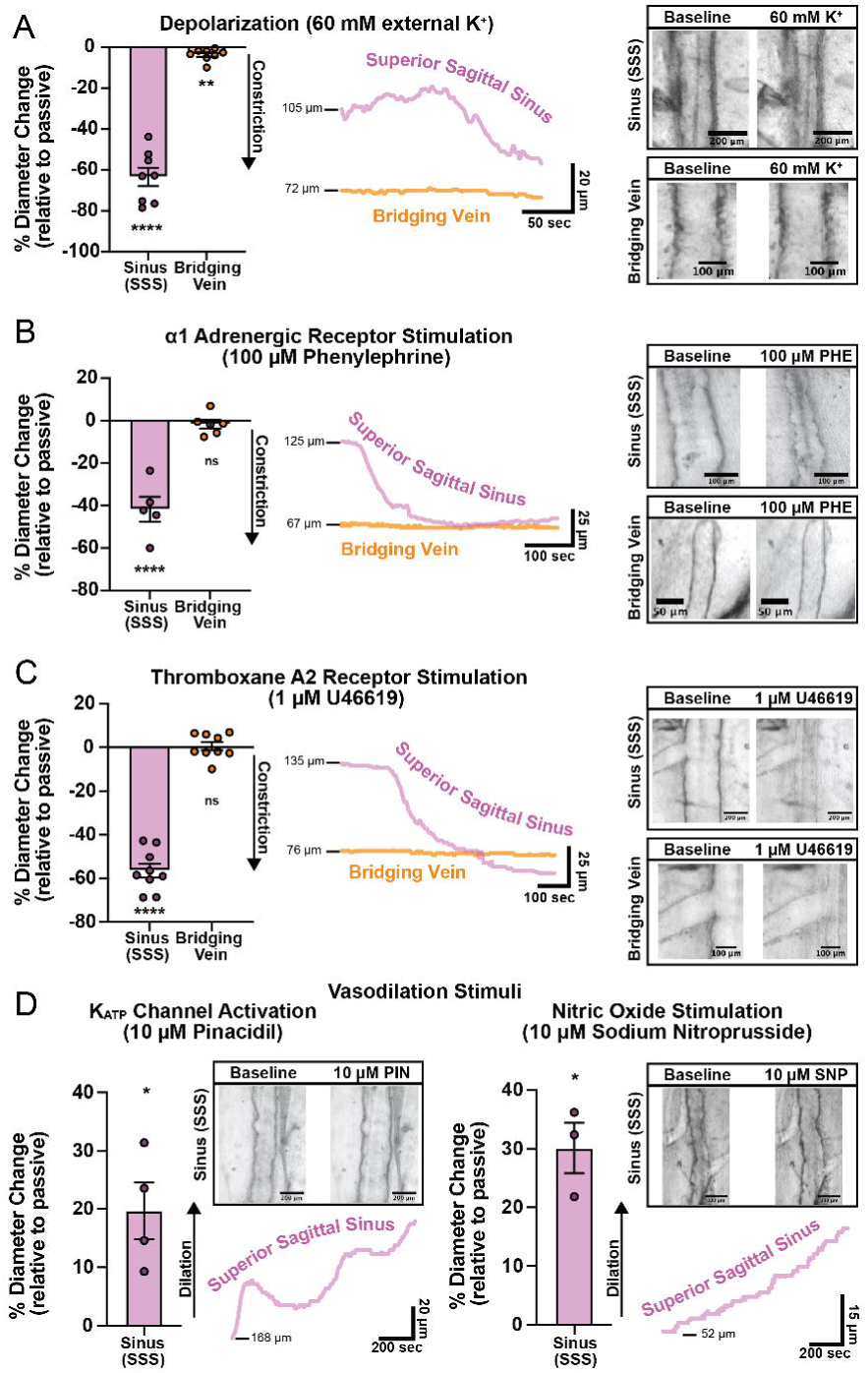
Dural venous sinuses constrict and dilate in response to smooth muscle contractile and relaxation stimuli. (A) Depolarization-induced constriction of the superior sagittal sinus (SSS) and attached bridging veins in response to 60 mM extracellular K⁺. Summarized constriction data are shown together with representative diameter traces over time and representative pre- and post-stimulus images of the SSS and bridging vein. Depolarization produced robust constriction of the SSS and only minimal constriction of attached bridging veins. (B) α_1_-adrenergic receptor stimulation with 100 µM phenylephrine produces robust constriction of the SSS but not attached bridging veins. Summarized constriction data are shown together with representative diameter traces over time and representative pre- and post-stimulus images of the SSS and bridging vein. (C) Thromboxane A2 receptor stimulation with 1 µM U46619 produces robust constriction of the SSS but not attached bridging veins. Summarized constriction data are shown together with representative diameter traces over time and representative pre- and post-stimulus images of the SSS and bridging vein. (D) Relaxation responses of preconstricted dural venous sinuses to ATP-sensitive potassium channel activation and nitric oxide exposure. Application of 10 µM pinacidil dilated the preconstricted SSS, and application of 10 µM sodium nitroprusside (SNP), a nitric oxide donor, produced additional sinus dilation. Summarized dilation data are shown together with representative SSS diameter traces over time and representative pre- and post-stimulus images of the SSS. Individual values are shown with mean ± SEM; n = 3–9 preparations. Statistical significance was determined by Wilcoxon signed-rank test versus a hypothetical value of zero, representing no constriction or dilation. *p < 0.05, **p < 0.01, ***p < 0.005, ****p < 0.001; ns, not significant.

To examine the vasodilatory capacity of sinus SMCs, pressurized sinus preparations were preconstricted with 5 mmHg pressure and 1 µM U46619. We tested dilation responses to activation of hyperpolarizing ATP-sensitive potassium (K_ATP_) channels or nitric oxide (NO) exposure. Application of 10 uM pinacidil, a synthetic K_ATP_ channel opener, led to 19.7 ± 4.9% dilation and in response to 10 uM sodium nitroprusside (SNP), an NO donor, sinus vessels dilated 30.1 ± 4.3% (Fig. 7 D,E). Movies 11 and 12 demonstrate sinus relaxation in response to pinacidil and SNP, respectively. Together, these results demonstrate that dural venous sinus SMCs bidirectionally regulate vessel diameter through both contractile and relaxation pathways. These findings establish the dural venous sinus as an actively regulated contractile segment of the cerebral venous circulation.

## DISCUSSION

The dural venous sinuses are central conduits for cerebral venous drainage and cerebrospinal fluid outflow, yet their potential for active diameter regulation has remained poorly defined. Sinus narrowing is often interpreted through the lens of passive mechanics, extrinsic compression, or structural remodeling [5, 6]. However, clinical observations that sinus caliber can change with altered cerebrospinal fluid pressure, together with the recurrence of stenosis adjacent to stented segments, suggest that sinus diameter may also be dynamically regulated [11, 13] [14]. Recent in vivo imaging further supports the idea that the superior sagittal sinus can undergo diameter changes over physiologically relevant timescales, including in response to behavior-linked changes in intracranial pressure and modulation of CGRP signaling[16] [15]). These observations raise a fundamental mechanistic question: whether the dural venous sinuses contain specialized mural cells capable of actively regulating venous tone. Here, we identify a distinct population of artery-like smooth muscle cells within dural venous sinuses and demonstrate that these cells confer robust contractile and dilatory control over sinus diameter.

Reanalysis of published single-cell RNA-sequencing data from mouse dura mater revealed molecular heterogeneity among dural mural cells [21]. Subclustering resolved three smooth muscle cell populations and a pericyte population, including an arterial-like SMC cluster enriched for canonical contractile genes, a venous-like population enriched for venous markers, and a third population with a combined venous and contractile transcriptional profile. This hybrid molecular identity was marked by expression of venous-associated genes together with contractile SMC markers, suggesting the presence of mural cells that are venous in vascular context but artery-like in contractile phenotype. Although the number of mural cells in this dataset was limited, the resolution was sufficient to identify candidate markers for spatial validation. Future studies using sinus-enriched SMC isolation and targeted single-cell or single-nucleus profiling will be important for defining this population more comprehensively, including its developmental origin, molecular diversity, and relationship to arterial and venous mural cell states.

Immunohistochemical analysis localized this contractile mural cell phenotype specifically to the dural venous sinuses. In whole-mount mouse dura, ACTA2 and CNN1 were strongly expressed by elongated SMCs surrounding the superior sagittal and transverse sinuses, whereas these markers were rapidly lost at the transition into bridging veins. This anatomical restriction was striking: bridging veins and non-sinus dural veins lacked the same contractile marker expression, while dural arteries and pial arteries retained the expected arterial SMC labeling. Importantly, this pattern was conserved in human dura mater, where ACTA2-positive SMCs lined the superior sagittal sinus but were largely absent from adjacent bridging veins. Together, these findings demonstrate that contractile SMCs are not broadly distributed throughout the dural venous network but are instead selectively concentrated around the major sinus vessels.

We next asked whether these artery-like sinus SMCs possess functional signaling properties associated with contractile vascular smooth muscle. Using live dural tissue from Myh11-GCaMP6f mice, we observed abundant localized subcellular Ca^2+^ events in sinus SMCs at baseline. Increasing intraluminal pressure from 0 to 5 mmHg increased the frequency, amplitude, and duration of these discrete Ca^2+^ signals. In arterial SMCs, localized Ca^2+^ events, including Ca^2+^ sparks, are coupled to membrane potential and vascular tone through activation of nearby potassium channels, forming a feedback mechanism that limits excessive constriction [26, 27, 31, 32]. Although the molecular identity and downstream targets of the sinus SMC Ca^2+^ events remain to be established, their pressure sensitivity indicates that sinus SMCs actively detect and respond to changes in transmural pressure. These findings place sinus SMCs in functional alignment with excitable contractile mural cells rather than passive venous wall elements.

Consistent with this pressure-sensitive Ca^2+^ signaling, dural venous sinuses constricted robustly in response to increased intraluminal pressure. Using an ex vivo pressurized sinus preparation, we found that the superior sagittal sinus narrowed across a graded range of pressures, with prominent constriction at physiologically relevant pressures, whereas adjacent bridging veins showed little or no comparable response. The pressure-induced decrease in sinus diameter was Ca^2+^ sensitive, as removal of extracellular Ca^2+^ increased passive diameter and diminished active tone. This behavior is consistent with pressure-dependent contractile mechanisms described in resistance arteries and other contractile vascular beds [28, 29]. These data support a model in which changes in sinus caliber are not explained solely by passive collapse or deformation but instead reflect active, Ca^2+^ -dependent contractility of sinus SMCs. This distinction is important because active sinus tone provides a cellular mechanism through which venous outflow resistance could be rapidly adjusted in response to changes in intracranial pressure, cerebral blood volume, or downstream drainage conditions.

Sinus SMCs also responded to canonical contractile stimuli associated with arterial smooth muscle excitability. Pressure, membrane depolarization, thromboxane A2 receptor activation, α_1_-adrenergic receptor stimulation, and electrical field stimulation each produced robust global increases in sinus SMC Ca^2+^. These responses indicate that sinus SMCs engage major signaling pathways classically associated with contractile vascular smooth muscle, including voltage-dependent Ca^2+^ entry and Gq-coupled receptor signaling [27, 33–36]. The α_1_-adrenergic responsiveness is particularly notable given the dense sympathetic innervation of the superior sagittal sinus, consistent with prior descriptions of adrenergic innervation in vascular beds and venous structures [17, 37]. Electrical field stimulation increased sinus SMC Ca^2+^, and this response was strongly reduced by α_1_-adrenergic receptor blockade, indicating that endogenous sympathetic nerve activity can directly engage sinus SMC Ca^2+^ signaling. Thus, sinus SMCs are positioned to integrate both mechanical and neural inputs.

These Ca^2+^ responses translated into strong changes in vessel diameter. Depolarization, α_1_-adrenergic receptor activation, and thromboxane A2 receptor stimulation each produced robust constriction of the superior sagittal sinus. In contrast, bridging veins showed only minimal constriction to depolarization and no detectable constriction to phenylephrine or U46619. This functional divergence mirrors the anatomical restriction of contractile SMC markers to the sinus and indicates that the dural venous system is regionally specialized: large sinus vessels are equipped for active tone generation, whereas bridging veins appear comparatively non-contractile under the conditions tested here. This specialization may allow the sinus to serve as a regulated outflow segment within the cerebral venous circulation while preserving distinct mechanical properties in upstream collecting veins.

Sinus SMCs also retained the capacity for active relaxation. Pinacidil dilated preconstricted sinuses, indicating that K_ATP_ channel activation can oppose sinus tone, while sodium nitroprusside produced robust dilation, consistent with nitric oxide-sensitive relaxation pathways. These responses align with established mechanisms by which membrane hyperpolarization and nitric oxide signaling reduce vascular smooth muscle tone [20, 38–40]. Together with the constrictor responses described above, these findings demonstrate that dural venous sinus diameter is bidirectionally regulated rather than passively determined by pressure alone.

This active control has important implications for cerebral venous outflow. Because the dural venous sinuses sit at the convergence of cerebral blood drainage and CSF outflow, even modest changes in sinus caliber could influence venous pressure, cerebral blood volume, CSF drainage, and intracranial pressure [3, 5, 7, 9, 10, 12, 23, 41]. Sympathetic regulation provides one mechanism by which sinus tone could be adjusted during changes in behavioral state, systemic cardiovascular demand, or intracranial pressure. Conversely, sustained or inappropriate sinus constriction could impair venous drainage and contribute to elevated venous pressure gradients, a clinically relevant feature of venous sinus stenosis [11, 13, 14]. Thus, these findings support a model in which dural venous sinuses function not only as drainage conduits, but as actively regulated vascular structures capable of influencing brain-wide hemodynamics and fluid clearance.

Several limitations should guide future work. First, although our data establish that sinus SMCs are contractile and pressure sensitive ex vivo, the extent to which these mechanisms regulate sinus diameter in vivo remains to be determined. Future experiments combining in vivo sinus imaging with targeted manipulation of SMC excitability, sympathetic signaling, or specific ion channels will be required to define how these mechanisms operate under physiological conditions. Second, we did not assess sex-dependent differences in sinus SMC structure or function. This is an important direction given the clinical association between venous sinus stenosis and disorders such as idiopathic intracranial hypertension, which disproportionately affect women of reproductive age [42–44]. Third, the present study focused primarily on SMC-intrinsic mechanisms and sympathetic input. Endothelial cells, sensory fibers, parasympathetic or other autonomic inputs, local inflammatory mediators, and paracrine signaling within the dura may all modulate sinus tone. Defining these interactions will be essential for understanding how sinus diameter is controlled in normal physiology and how it may become dysregulated in disease.

In summary, this study identifies the dural venous sinus as a specialized contractile segment of the cerebral venous circulation. Sinus vessels contain a distinct population of artery-like SMCs with venous molecular identity, arterial contractile protein expression, pressure-sensitive Ca^2+^ signaling, myogenic constriction, adrenergic responsiveness, and bidirectional control of vessel diameter. These findings reveal a previously underappreciated mechanism for active regulation of cerebral venous outflow and suggest that sinus SMCs may play an important role in coordinating cerebral blood drainage, cerebrospinal fluid clearance, and intracranial pressure homeostasis.

## METHODS

### Animals-

Animal use and procedures were in accordance with protocols approved by the University of Vermont Institutional Animal Care and Use Committee (IACUC). Dural and brain tissues were taken from 2-4 month old male and female C57bl/6J mice (stock no. 000664; Jackson Laboratories). Tamoxifen-inducible Myh11-GCaMP6f mice (4-6 months old male and female) were used to observe Ca^2+^ signals in venous SMCs. These mice were generated by crossing cre-dependent GCaMP6f mice (stock no. 028865; Jackson Laboratories) with Myh11-Cre-ERT2-RAD (stock no. 037658; Jackson Laboratories) mice. Tamoxifen dissolved in safflower oil (40 mg/mL) was administered via intraperitoneal injection to 4–8-week old mice with 5 consecutive 100 mg/kg daily doses. All animals were group housed with an enriched environment on a 12-hour light/dark cycle with free access to food and water. Mice were euthanized by 5% isoflurane overdose or intraperitoneal injection of pentobarbital (100mg/kg), followed by rapid decapitation.

### Reanalysis of dura mater single cell RNA sequencing data set-

Mural cell molecular characterization was achieved using previously published single cell RNA sequencing data [21]

### Immunohistochemistry-

Mice were euthanized via 5% isoflurane and perfused via transcranial perfusion with ice-cold phosphate buffered saline with 1% heparin followed by ice-cold 4% paraformaldehyde (PFA). Calvaria and brain were removed and further fixed in 4% PFA at 4 °C for 2 hours. The skull cap and brain were removed and thoroughly washed with PBS. The whole dura attached to the calvaria was carefully dissected and placed in PBS. Using a vibratome, brain tissue was sectioned into 200 um slices. Slices included coronal sections and sections along the pial surface. The tissues were blocked overnight at 4 °C in blocking buffer (PBS with 1% bovine serum albumin (BSA), 0.5% Triton X-100, 2.5% donkey serum). Tissues were then incubated overnight at 4 °C in antibodies or stains dissolved in blocking buffer: anti-α smooth muscle actin (1:500, ACTA2, Sigma catalog F3777), anti-CD31/PECAM-1 (1:500, R&D systems catalog AF3628), anti-calponin 1 (1:500, CNN1, Abcam catalog AB46794). Tissue was further washed and stained with species specific fluorescently conjugated secondary antibodies (Thermo Fisher Alex Fluor Plus Secondaries) in blocking buffer at room temperature for 2 hr. Dura tissue and brain slices were mounted on a slide using Vactashied mounting medium. Some dura tissue was pinned flat on a silicone bottom Petri dish with PBS medium and imaged before mounting on slides.

### Pressurized and flat mount sinus preparation-

The top of the skull (whole calvaria) was excised from euthanized mice and placed in ice-cold HEPES dissecting solution consisting of 130 mM NaCl, 3 mM KCl, 2 mM CaCl_2_, 1 mM MgCl_2_, 4 mM glucose, and 10 mM HEPES at pH 7.4. The dura was carefully peeled off using fine forceps. The ex vivo dural preparation was transferred and pinned within an adapted pressure myography recording chamber with a silicone platform (IMF, UVM) containing cold bicarbonate-buffered physiological saline solution (PSS) consisting of 119 mM NaCl, 3 mM KCl, 1 mM MgCl_2_, 2 mM CaCl_2_, 26.2 mM NaHCO_3_, 1 mM NaPO_4_, and 5 mM glucose; bubbled with 95%O_2_/5% CO_2_ gas (pH ∼7.35). The dural tissue was pinned flat. ventral side up, using 70 um tungsten wire pins. A borosilicate glass cannula was pulled with tip opening of ∼200 um (Sutter Instruments, Novato, CA), fixed into the chamber-attached micromanipulator, and backfilled with PSS. The glass cannula was inserted into the superior sagittal sinus and tied with fine nylon thread. The chamber was then transferred to the imaging microscope and perfused with bubbled PSS at a rate of 3 mL/min and a bath temperature of ∼36C. The cannula was attached to a gravity column containing PSS to deliver specific intraluminal pressures. Pressure values were verified using an in-line pressure transducer and meter (PM-4, Living Systems Instrumentation). Passive diameters were achieved using zero calcium conditions, zero calcium PSS was made by removing CaCl_2_, and increasing MgCl_2_ concentration to 3 mM (to maintain external divalent concentration) and adding 5 mM EGTA.

To detect pressure-induced contractility in the pressurized sinus preparation, changes in vessel diameter were assessed after 15 minutes of pressure, agonist, or depolarizing solutions. Passive diameters were assessed at each pressure following 30 mins of zero calcium conditions.

Pressure-induced tone was assessed using the following equation:

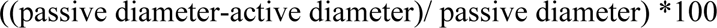

Preconstricted pressurized sinus preparations were exposed to pinacidil or SNP to test hyperpolarizing dilation effect in dural sinus SMCs. Dilation was assessed using the following equation:

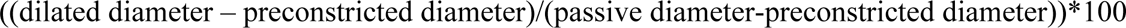

Changes in GCaMP6f fluorescence levels were used to assess pressure, agonist, and depolarizing effects on sinus SMC cytosolic Ca^2+^ levels. For agonist and depolarizing conditions flat mount sinus preparations were used as described above without cannulation. For pressure-induced Ca^2+^ changes sinus preparations were used as described above. For global cytosolic GCaMP fluorescent changes, the baseline whole FOV GCaMP6f signal was compared to the whole FOV GCaMP6f signal during and following stimuli (as previously demonstrated [33]). Localized Ca^2+^ event was achieved using SparkAn software as previously described [45], briefly subcellular Ca^2+^ event criteria included an increased F/F_0_ amplitude >1.1 with baseline and event fluorescent intensity >2 standard deviations.

Electrical Field Stimulation: Stimulation of sympathetic nerve fibers was achieved with sinus bordering platinum electrodes attached to Grass Stimulator. Frequency and pulse width were biased to sympathetic nerve activation [46] and were accordingly set at 10-90V, 0.25 ms pulse width, and 20 Hz frequency.

### Widefield and Confocal Fluorescent Microscopy -

Fixed tissues were imaged using widefield epifluorescent microscopy (Olympus MVX10, Andor iXon Ultra 888, X-Cite Xylis LED) or spinning disk confocal microscopy (Upright Nikon Ti-E, Yokogawa CSU-W, Dual Andor iXon Ultra 897, Andor Laser Combiner – 488, 546, 647 nm emission wavelengths, Nikon Elements Software).

An upright Olympus BX51W microscope with near infrared optics and attached CMOS camera was used to image sinus edges. Diameter measurements were made using ImageJ analysis software and customized macros to assess real-time edge to edge distance.

GCaMP6f fluorescence was detected in live tissue using the spinning disk confocal microscope described above.

## Acknowledgements

We thank Todd Clason for microscopy assistance. This study was supported by grants from the Totman Medical Research Trust, The Cardiovascular Research Institute of Vermont, The American Heart Association 23CDA1050558, and The National Heart Lung and Blood Institute of the National Institutes of Health K01HL167052 to N.R.K. National Institute of General Medical Sciences of the National Institutes of Health P20GM135007 Pilot Project Grant and Custom Physiology and Imaging Core resources to N.R.K and A.S.S.B. Data and effort provided by L.H. and C.B. was not supported by the funding listed above. Collaborative dialog and efforts between C.B. and N.R.K were facilitated by the Leducq Foundation Transatlantic Network of Excellence (International Network of Excellence on Brain Endothelium: A Nexus for Cerebral Small Vessel Disease 22CVD01 BRENDA.

## Contributions

N.R.K. supervised study design, data acquisition, data analysis, and manuscript preparation. N.R.K and A.S.S.B conceptualized the project. N.R.K., M.E.L, H.C.R, and L.S.H designed experiments, collected data, and analyzed data. J.C.D. prepared and optimized embedded human dura mater samples. L.H. and C.B. generated original RNA-sequencing data and reanalyzed mural cell clusters. N.R.K and M.E.L wrote the manuscript.

## Movies and Movie Legends

Movie 1. Localized Ca^2+^ events in dural venous sinus smooth muscle cells.

Live whole-mount dura mater from Myh11-GCaMP6f mice was used to visualize Ca^2+^ activity in dural venous sinus smooth muscle cells from an ex vivo pressurized sinus preparation. GCaMP6f fluorescence was recorded by spinning disk confocal microscopy. Left: raw GCaMP6f fluorescence displayed with a green lookup table. Middle: raw fluorescence over time, showing distinct localized subcellular Ca^2+^ events in sinus smooth muscle cells. Right: extracted Ca^2+^ events identified using event detection criteria, with event amplitude displayed as F/F_0_ fluorescence changes.

Movie 2. Pressure-induced constriction of the superior sagittal sinus.

Live whole-mount dura mater was used in an ex vivo pressurized sinus preparation to visualize diameter changes in the superior sagittal sinus. The superior sagittal sinus was cannulated, bath perfused, and imaged by near-infrared widefield microscopy using a 4X, 0.10 NA objective. The movie shows constriction of the superior sagittal sinus during an increase in intraluminal pressure from 0 to 5 mmHg over a 1194 s recording period.

Movie 3. Pressure-evoked global Ca^2+^ increase in dural venous sinus smooth muscle cells. Live whole-mount dura mater from Myh11-GCaMP6f mice was imaged by spinning disk confocal microscopy to visualize GCaMP6f fluorescence in dural venous sinus smooth muscle cells. Images were acquired with a 100 ms exposure and displayed using a fire lookup table, where low fluorescence intensity appears black to dark blue and high fluorescence intensity appears red to white. Movie shows the smooth muscle Ca^2+^ response during increased intraluminal pressure from 0 to 5 mmHg.

Movie 4. Depolarization-evoked global Ca^2+^ increase in dural venous sinus smooth muscle cells. Live whole-mount dura mater from Myh11-GCaMP6f mice was imaged by spinning disk confocal microscopy to visualize GCaMP6f fluorescence in dural venous sinus smooth muscle cells. Images were acquired with a 100 ms exposure and displayed using a fire lookup table, where low fluorescence intensity appears black to dark blue and high fluorescence intensity appears red to white. Movie shows the smooth muscle Ca^2+^ response to membrane depolarization with 60 mM extracellular K⁺.

Movie 5. Thromboxane A2 receptor activation evokes a global Ca^2+^ increase in dural venous sinus smooth muscle cells.

Live whole-mount dura mater from Myh11-GCaMP6f mice was imaged by spinning disk confocal microscopy to visualize GCaMP6f fluorescence in dural venous sinus smooth muscle cells. Images were acquired with a 100 ms exposure and displayed using a fire lookup table, where low fluorescence intensity appears black to dark blue and high fluorescence intensity appears red to white. Movie shows the smooth muscle Ca^2+^ response to thromboxane A2 receptor activation with 1 µM U46619.

Movie 6. α_1_-adrenergic receptor stimulation evokes a global Ca^2+^ increase in dural venous sinus smooth muscle cells.

Live whole-mount dura mater from Myh11-GCaMP6f mice was imaged by spinning disk confocal microscopy to visualize GCaMP6f fluorescence in dural venous sinus smooth muscle cells. Images were acquired with a 100 ms exposure and displayed using a fire lookup table, where low fluorescence intensity appears black to dark blue and high fluorescence intensity appears red to white. Movie shows the smooth muscle Ca^2+^ response to α_1_-adrenergic receptor stimulation with 100 µM phenylephrine.

Movie 7. Electrical field stimulation evokes a global Ca^2+^ increase in dural venous sinus smooth muscle cells.

Live whole-mount dura mater from Myh11-GCaMP6f mice was imaged by spinning disk confocal microscopy to visualize GCaMP6f fluorescence in dural venous sinus smooth muscle cells. Images were acquired with a 100 ms exposure and displayed using a fire lookup table, where low fluorescence intensity appears black to dark blue and high fluorescence intensity appears red to white. Movie shows the smooth muscle Ca^2+^ response to electrical field stimulation of perivascular nerves.

Movie 8. Depolarization-induced constriction of the superior sagittal sinus.

Live whole-mount dura mater was used in an ex vivo pressurized sinus preparation to visualize stimulus-induced changes in vessel diameter. The superior sagittal sinus was cannulated, bath perfused, and maintained at 5 mmHg intraluminal pressure. Vessel walls were visualized by near-infrared widefield microscopy using a 4X, 0.10 NA objective to track sinus diameter over time. The movie shows constriction of the superior sagittal sinus in response to membrane depolarization with 60 mM extracellular K⁺ over a 199 s recording period.

Movie 9. α_1_-adrenergic receptor stimulation constricts the superior sagittal sinus.

Live whole-mount dura mater was used in an ex vivo pressurized sinus preparation to visualize stimulus-induced changes in vessel diameter. The superior sagittal sinus was cannulated, bath perfused, and maintained at 5 mmHg intraluminal pressure. Vessel walls were visualized by near-infrared widefield microscopy using a 4X, 0.10 NA objective to track sinus diameter over time. The movie shows constriction of the superior sagittal sinus in response to α_1_-adrenergic receptor stimulation with 100 µM phenylephrine over a 468 s recording period.

Movie 10. Thromboxane A2 receptor stimulation constricts the superior sagittal sinus.

Live whole-mount dura mater was used in an ex vivo pressurized sinus preparation to visualize stimulus-induced changes in vessel diameter. The superior sagittal sinus was cannulated, bath perfused, and maintained at 5 mmHg intraluminal pressure. Vessel walls were visualized by near-infrared widefield microscopy using a 4X, 0.10 NA objective to track sinus diameter over time. The movie shows constriction of the superior sagittal sinus in response to thromboxane A2 receptor stimulation with 1 µM U46619 over a 580 s recording period.

Movie 11. ATP-sensitive potassium channel activation dilates the preconstricted superior sagittal sinus.

Live whole-mount dura mater was used in an ex vivo pressurized sinus preparation to visualize stimulus-induced changes in vessel diameter. The superior sagittal sinus was cannulated, bath perfused, maintained at 5 mmHg intraluminal pressure, and preconstricted with 1 µM U46619. Vessel walls were visualized by near-infrared widefield microscopy using a 4X, 0.10 NA

objective to track sinus diameter over time. The movie shows dilation of the preconstricted superior sagittal sinus in response to ATP-sensitive potassium channel activation with 10 µM pinacidil over a 1048 s recording period.

Movie 12. Nitric oxide donor application dilates the preconstricted superior sagittal sinus.

Live whole-mount dura mater was used in an ex vivo pressurized sinus preparation to visualize stimulus-induced changes in vessel diameter. The superior sagittal sinus was cannulated, bath perfused, maintained at 5 mmHg intraluminal pressure, and preconstricted with 1 µM U46619. Vessel walls were visualized by near-infrared widefield microscopy using a 4X, 0.10 NA objective to track sinus diameter over time. The movie shows dilation of the preconstricted superior sagittal sinus in response to the nitric oxide donor sodium nitroprusside (10 µM) over a 1048 s recording period.

## REFERENCES

1. Balo, J., The dural venous sinuses. Anat Rec, 1950. 106(3): p. 319–24.

2. Retzius, A.K.a.M.G., Studien in der Anatomie des Nervensystems und des Bindegewebes. Vol. 2. 1875–1876: Samson & Wallin.

3. Schaeffer, S. and C. Iadecola, Revisiting the neurovascular unit. Nat Neurosci, 2021. 24(9): p. 1198–1209.

4. Melnikow-Raswedenkow, N., Histologische Untersuchungen über den normalen Bau der Dura mater und über Pachymeningitis interna. Beiträge zur pathologischen Anatomie und zur allgemeinen Pathologie, 1890. 28: p. 217–254.

5. Proulx, S.T., Cerebrospinal fluid outflow: a review of the historical and contemporary evidence for arachnoid villi, perineural routes, and dural lymphatics. Cell Mol Life Sci, 2021. 78(6): p. 2429–2457.

6. Sundararajan, S.H., et al., Dural Venous Sinus Stenosis: Why Distinguishing Intrinsic-versus-Extrinsic Stenosis Matters. AJNR Am J Neuroradiol, 2021. 42(2): p. 288–296.

7. Friedman, D.I., Cerebral venous pressure, intra-abdominal pressure, and dural venous sinus stenting in idiopathic intracranial hypertension. J Neuroophthalmol, 2006. 26(1): p. 61–4.

8. Lawton, M.T., R. Jacobowitz, and R.F. Spetzler, Redefined role of angiogenesis in the pathogenesis of dural arteriovenous malformations. J Neurosurg, 1997. 87(2): p. 267–74.

9. Jones, H.C. and J.A. Gratton, The effect of cerebrospinal fluid pressure on dural venous pressure in young rats. J Neurosurg, 1989. 71(1): p. 119–23.

10. Gurney, S.P., et al., Exploring The Current Management Idiopathic Intracranial Hypertension, And Understanding The Role Of Dural Venous Sinus Stenting. Eye Brain, 2020. 12: p. 1–13.

11. Fargen, K.M., et al., “Idiopathic” intracranial hypertension: An update from neurointerventional research for clinicians. Cephalalgia, 2023. 43(4): p. 3331024231161323.

12. Shukla, V., L.A. Hayman, and K.H. Taber, Adult cranial dura II: venous sinuses and their extrameningeal contributions. J Comput Assist Tomogr, 2003. 27(1): p. 98–102.

13. Lim, J., et al., Stenting for Venous Sinus Stenosis in Patients With Idiopathic Intracranial Hypertension: An Updated Systematic Review and Meta-Analysis of the Literature. Neurosurgery, 2024. 94(4): p. 648–656.

14. Fargen, K.M., A unifying theory explaining venous sinus stenosis and recurrent stenosis following venous sinus stenting in patients with idiopathic intracranial hypertension. J Neurointerv Surg, 2021. 13(7): p. 587–592.

15. Zhang, Q., et al., Ultrafast venous and sagittal sinus constrictions in the brain driven by abdominal pressure. bioRxiv, 2026: p. 2026.05.02.722426.

16. Monaghan, K.L., et al., Highly dynamic dural sinuses support meningeal immunity. Nature, 2026.

17. Furness, J.B., The adrenergic innervation of the vessels supplying and draining the gastrointestinal tract. Z Zellforsch Mikrosk Anat, 1971. 113(1): p. 67–82.

18. Chihara, E., et al., Comparative effects of nitroglycerin on intestinal vascular capacitance and conductance. Can J Cardiol, 2002. 18(2): p. 165–74.

19. Shi, W., D.H. Eidelman, and R.P. Michel, Differential relaxant responses of pulmonary arteries and veins in lung explants of guinea pigs. J Appl Physiol (1985), 1997. 83(5): p. 1476–81.

20. Barnes, P.J. and S.F. Liu, Regulation of pulmonary vascular tone. Pharmacol Rev, 1995. 47(1): p. 87–131.

21. Pietila, R., et al., Molecular anatomy of adult mouse leptomeninges. Neuron, 2023. 111(23): p. 3745–3764 e7.

22. Arroyo-Ataz, G., et al., Single-Cell Transcriptomics and Lineage Tracing Unveil Parallels in Lymphatic Muscle and Venous Smooth Muscle Development, Identity, and Function. Arterioscler Thromb Vasc Biol, 2025. 45(7): p. 1207–1225.

23. Betsholtz, C., et al., Advances and controversies in meningeal biology. Nat Neurosci, 2024. 27(11): p. 2056–2072.

24. Ushiwata, I. and T. Ushiki, Cytoarchitecture of the smooth muscles and pericytes of rat cerebral blood vessels. A scanning electron microscopic study. J Neurosurg, 1990. 73(1): p. 82–90.

25. Vanlandewijck, M., et al., A molecular atlas of cell types and zonation in the brain vasculature. Nature, 2018. 554(7693): p. 475–480.

26. Amberg, G.C. and M.F. Navedo, Calcium dynamics in vascular smooth muscle. Microcirculation, 2013. 20(4): p. 281–9.

27. Nelson, M.T., et al., Calcium channels, potassium channels, and voltage dependence of arterial smooth muscle tone. Am J Physiol, 1990. 259(1 Pt 1): p. C3–18.

28. Knot, H.J. and M.T. Nelson, Regulation of arterial diameter and wall [Ca2+] in cerebral arteries of rat by membrane potential and intravascular pressure. J Physiol, 1998. 508 (**Pt 1**)(Pt 1): p. 199–209.

29. Davis, M.J., et al., Myogenic constriction and dilation of isolated lymphatic vessels. Am J Physiol Heart Circ Physiol, 2009. 296(2): p. H293–302.

30. Keller, J.T., et al., Sympathetic innervation of the supratentorial dura mater of the rat. The Journal of Comparative Neurology, 1989. 290(2): p. 310–321.

31. Nelson, M.T., et al., Relaxation of arterial smooth muscle by calcium sparks. Science, 1995. 270(5236): p. 633–7.

32. Jaggar, J.H., et al., Calcium sparks in smooth muscle. Am J Physiol Cell Physiol, 2000. 278(2): p. C235–56.

33. Klug, N.R., et al., Intraluminal pressure elevates intracellular calcium and contracts CNS pericytes: Role of voltage-dependent calcium channels. Proc Natl Acad Sci U S A, 2023. 120(9): p. e2216421120.

34. Pires, P.W., et al., The angiotensin II receptor type 1b is the primary sensor of intraluminal pressure in cerebral artery smooth muscle cells. J Physiol, 2017. 595(14): p. 4735–4753.

35. Pietrobon, D. and P. Hess, Novel mechanism of voltage-dependent gating in L-type calcium channels. Nature, 1990. 346(6285): p. 651–5.

36. Catterall, W.A., Voltage-gated calcium channels. Cold Spring Harb Perspect Biol, 2011. 3(8): p. a003947.

37. Bruno, R.M., et al., Sympathetic regulation of vascular function in health and disease. Front Physiol, 2012. 3: p. 284.

38. Standen, N.B., et al., Hyperpolarizing vasodilators activate ATP-sensitive K+ channels in arterial smooth muscle. Science, 1989. 245(4914): p. 177–80.

39. Robertson, B.E., et al., cGMP-dependent protein kinase activates Ca-activated K channels in cerebral artery smooth muscle cells. Am J Physiol, 1993. 265(1 Pt 1): p. C299–303.

40. Tare, M., et al., Hyperpolarization and relaxation of arterial smooth muscle caused by nitric oxide derived from the endothelium. Nature, 1990. 346(6279): p. 69–71.

41. Liu, K.C., et al., Venous sinus stenting for reduction of intracranial pressure in IIH: a prospective pilot study. J Neurosurg, 2017. 127(5): p. 1126–1133.

42. Raz, E., et al., Emergence of Venous Stenosis as the Dominant Cause of Pulsatile Tinnitus. Stroke Vasc Interv Neurol, 2022. 2(4): p. e000154.

43. Daniels, A.B., et al., Profiles of obesity, weight gain, and quality of life in idiopathic intracranial hypertension (pseudotumor cerebri). Am J Ophthalmol, 2007. 143(4): p. 635–41.

44. Ko, M.W., et al., Weight gain and recurrence in idiopathic intracranial hypertension: a case-control study. Neurology, 2011. 76(18): p. 1564–7.

45. Dabertrand, F., M.T. Nelson, and J.E. Brayden, Acidosis dilates brain parenchymal arterioles by conversion of calcium waves to sparks to activate BK channels. Circ Res, 2012. 110(2): p. 285–94.

46. Nausch, L.W., et al., Sympathetic nerve stimulation induces local endothelial Ca2+ signals to oppose vasoconstriction of mouse mesenteric arteries. Am J Physiol Heart Circ Physiol, 2012. 302(3): p. H594–602.

